# APOE4 carriers resistant to cognitive decline show unique relationships between cerebrovascular response to exercise and dual-task cognitive-balance performance

**DOI:** 10.1101/2025.09.30.679544

**Authors:** Jacqueline A. Palmer, Sandra A. Billinger

**Affiliations:** Division of Physical Therapy and Rehabilitation Science, Medical School, University of Minnesota, Minneapolis, MN, United States of America; Department of Neurology, School of Medicine, University of Kansas Medical Center, Kansas City, KS, United States of America; University of Kansas Alzheimer’s Disease Research Center, Fairway, KS, United States of America; Department of Molecular & Integrative Physiology, University of Kansas Medical Center, Kansas City, KS, USA

## Abstract

**BACKGROUND:** Emerging research indicates cognitive dual-tasking is an early marker for cognitive impairment, with particular implications for *Apolipoprotein E4 (APOE4)* carriers who are at higher genetic risk for Alzheimer’s disease. While *APOE4* carriers typically show accelerated cognitive decline and impaired cerebrovascular function with aging, exceptions to this norm exist and may provide insights into resilience mechanisms. The relationship between cerebrovascular response and cognitive-motor dual-task performance in cognitively-normal *APOE4* carriers who maintain preserved function remains unclear.

**METHODS:** Thirty cognitively-normal older adults (76±4 years, 8 *APOE4* carriers, 22 noncarriers) completed clinical balance and cognitive testing under single-task and dual-task conditions. Balance performance was assessed as distance traversed during challenging beam walking. Cognitive performance was assessed as response time during an auditory Stroop test. Transcranial Doppler ultrasound measured cerebrovascular response to moderate-intensity aerobic exercise. We tested group differences in cognitive-balance dual task performance and relationships between cerebrovascular response and dual-task interference (DTI) in balance and cognitive domains, and effects of APOE4 genotype on these relationships.

**RESULTS:** No differences in cerebrovascular response or dual-task performance were observed between *APOE4* carriers and noncarriers. However, *APOE4* carriers displayed unique cerebrovascular-behavioral relationships. In *APOE4* carriers, higher cerebrovascular response to exercise was associated with less balance DTI (r=0.839, p=0.009) and less cognitive DTI (r=0.832, p=0.020), while no relationships were observed in noncarriers (p>0.187).

**CONCLUSIONS:** Cognitively-normal *APOE4* carriers with preserved cognitive-balance dual-task function represent exceptions that may model aging resilience mechanisms. The unique cerebrovascular-behavioral relationships suggest that maintaining cerebrovascular function provides preliminary support for neuromotor and neurocognitive resilience to dual-task challenges in genetically vulnerable populations.

## 1. Introduction

Approximately 25% of the U.S. population carry at least a single *Apolipoprotein E4* (*APOE4*) allele, the strongest known genetic risk factor for Alzheimer’s disease (AD)(1) and a high risk for other diseases involving cardiovascular health(e.g. heart attack and stroke).(2) Increasing evidence points to an early and key role of cerebrovascular dysfunction in the pathogenesis of Alzheimer’s disease.(3–5) The *APOE4* genotype has been the most commonly studied genetic variant linked to brain function and appears to act synergistically with cardiovascular health (e.g. blood pressure, lipid profiles, white matter hyperintensities) to influence cognitive decline.(2,6–9) When brain vascular function is attenuated, this can reduce or slow down the clearance of amyloid-beta – a neurobiological hallmark for AD- and promote its accumulation in the brain.(10) For *APOE4* carriers, who show accelerated amyloid-beta pathology,(11) maximizing cerebrovascular function may be critical for maintenance of brain health with aging. For example, cerebrovascular function can be characterized as responsivity under conditions of *physiologic stress* (e.g. sit-to-stand positional changes, aerobic exercise, heat stress, hypoxia)(12–18) that can play an important role in maintaining brain metabolism and function with aging,(19) as the damaging effects of repeated transient disruption of blood, glucose, and oxygen supply to brain tissue accumulate over time (20). Recent studies involving exercise interventions suggest that the protective effects of exercise on risk for future dementia appear to be even stronger in individuals who carry *APOE4* (21) (22). Supporting this notion our group previously showed that, despite having higher amyloid-beta deposition compared to noncarriers, cognitively-normal older adult *APOE4* carriers who maintained exceptionally high cerebrovascular response to a bout of aerobic exercise showed no difference in cognitive executive function performance (23). This is remarkable because of the high genetic vulnerability of APOE4 carriers to early signs of AD and the fact that impaired cognitive executive function is one of the earliest cognitive manifestations of mild cognitive impairment (MCI) that can progress to dementia (24,25).

Decline in balance and gait function may occur several years before individuals meet clinical diagnosis for mild cognitive impairment (MCI),(26–28) implicating that motor behavior may be a more sensitive indicator of underlying neuropathology that precipitates clinical cognitive syndrome. The emergence of cognitive interference in balance and walking over the course of aging is one of the most prevalent clinical phenomenon that emerges with aging (29,30), e.g. older adults cannot “walk and talk” at the same time (29). A person’s ability to perform cognitive-motor dual-tasking may reflect individual neural capacity (31–34) that supports neurocognitive and neuromotor resilience imposed by competing attentional demand (35). Clinical dual-task paradigms can be used to assess cognitive-motor interference, in which the individual is asked to simultaneously perform a cognitive task while balancing/walking and the change in their performance in either or both tasks is measured.(36,37) Specifically, greater degradation of *balance and gait* performance under cognitive loading (i.e. greater dual-task interference (DTI)) is an early and sensitive indicator of behavioral dysfunction in older adults,(29,38–43) and can predict future dementia(44) and falls.(38,45) One pilot study showed that older adult *APOE4* carriers showed greater cognitive DTI during walking, in which the cognitive task domain of executive function showed an even greater effect compared to a working memory task in *APOE4* carriers during dual-task gait.(36) Recently, our group showed that there is a relationship between higher cerebral blood velocity and cognitive-balance dual-task behavior in cognitively-normal older adults, especially with advanced age.(46) The effect of advanced age is notable because preserved cognitive function becomes more meaningful with age, as the magnitude of average decline in cognitive performance over time is disproportionately influenced by age, and manifestations of genetic risk becomes critical factors for disease risk.(47,48) This poses the question of whether modifiable factors such as brain vascular health contribute to the early clinical manifestations of cognitive-motor dual-task interference that could be therefore be targeted and modified with clinical intervention during preclinical disease stages in highly vulnerable older adults who carry *APOE4*.

Our group previously showed that, despite having greater amyloid-beta deposition, cognitively-normal *APOE4* carriers with more robust cerebrovascular response to aerobic exercise had higher cognitive response inhibition performance under single-task conditions.(23) Cerebrovascular response to physiologic stress like aerobic exercise, measurable through clinically feasible and cost-effective transcranial Doppler ultrasound (TCD),(17,49–51) serves as a more sensitive early indicator of cognitive dysfunction than resting assessments, which fail to detect subtle vascular impairments in older adults.(17,52) In this brief research report, we aimed to confirm and extend the pilot findings by Whitson et al (36) by assessing whether *APOE4* carriers who remained cognitively normal in the advanced stages of aging (70+ years) show lower single- and dual-task cognitive and dynamic balance performance compared to their noncarrier peers. We also explored whether our previous findings showing interactive effects between *APOE4* carriers and cerebrovascular response to exercise on single-task cognitive function(23) apply to cognitive-motor dual-task balance behavior.

## 2. Materials and Methods

### 2.1. Participants

Thirty participants (76±4 years, 19 females) from the University of Kansas Alzheimer’s Disease Research Center (P30AG072973)^3,10–12^ were selected for this study (**Table 1**). For inclusion in this analysis, participants had to be (1) age 70-90 years, (2) have normal cognition (see below), (3) have absence of neurologic or orthopedic disability to prevent independent standing and walking, and (4) speak the English language. Exclusion criteria were (1) insulin-dependent diabetes, (2) peripheral neuropathy affecting somatosensation, (3) active coronary artery disease and congestive heart failure. The University of Kansas Institutional Review Board approved this protocol (IRB#: STUDY 00147888) and all participants provided written informed consent.

**Table 1.** Participant characteristics.

| Table 1. Participant characteristics |  |  |  |  |
| --- | --- | --- | --- | --- |
|  | ALL | APOE4 | Noncarriers | P value |
|  | (n=30) | E4/E3 (n=8) | E3/E3 (n=22) |  |
| Age | 76 ± 4 | 75 ± 4 | 76 ± 5 | p=0.584 |
| Sex (F/M)<sup>∞</sup> | 19/11 | 3/5 | 16/6 | p=0.077 |
| Resting mean arterial pressure (mmHg) | 93 ± 22 | 96 ± 18 | 91 ± 24 | 0.585 |
| Resting Heart rate (bpm) | 73 ± 9 | 71 ± 11 | 74 ± 8 | 0.152 |
| Exercise heart rate (bpm) | 107 ±7 | 103 ± 10 | 109 ± 5 | p=0.056 |
| CBFv response to exercise | 1.6 ± 4.4 | 1.5 ± 3.84 | 1.65 ± 4.73 | p=0.943 |
| Exercise Watts | 50 ± 23 | 58 ± 32 | 47 ± 19 | p=0.266 |

Values are depicted as mean ± SD. \*p<0.05; <sup>∞</sup>self-reported

### 2.2. Clinical screening for cognitive impairment and eligibility

This exploratory investigation specifically aimed to compare cognitive-balance dual-task performance and relationships between cerebrovascular and cognitive-balance dual-task metrics between older adult *APOE4* carriers and noncarriers with well-preserved cognition. Participants completed extensive neuropsychological testing and Clinical Dementia Rating (CDR) assessment through the University of Kansas Alzheimer’s Disease Research Center, administered by a qualified clinician and psychometrist. Only individuals with normal cognitive status (CDR=0) and APOE genotyping were included in this analysis.

### 2.3. Behavioral assessments of balance and cognitive performance

Participants completed a challenging beam walking task to assess balance performance.(53–55) Participants walked across a 16-foot long narrow beam (3.5-inch width) (1-inch height to minimize postural threat).(53) Participants were instructed to walk as far as they could across the beam but did not receive instructions on walking speed. This dynamic beam walking balance task can reliably detect dynamic balance proficiency across older adults with high and low balance function.(54) This beam walk test can also detect age-related differences in cognitive-balance dual-task interference,(53) and may be more sensitive in detecting balance function compared to conventional clinical tests.(56) Older adults who achieved perfect beam scores on the first two trials were progressed to a narrower beam width (1-inch), and started on the narrow beam with a baseline score of 16 feet.

#### Single-task (ST) balance performance

Participants wore a safety belt and were instructed to fold their arms across their chest, fix their gaze straight ahead at a point on the wall straight ahead at eye level, and walk forward across the beam without stepping off the beam or uncrossing their arms. A trial was stopped when the participant stepped off the beam, walked sideways, or unfolded their arms, and their initial ground foot placement position was marked (**Figure 1A**).

**Figure 1.**
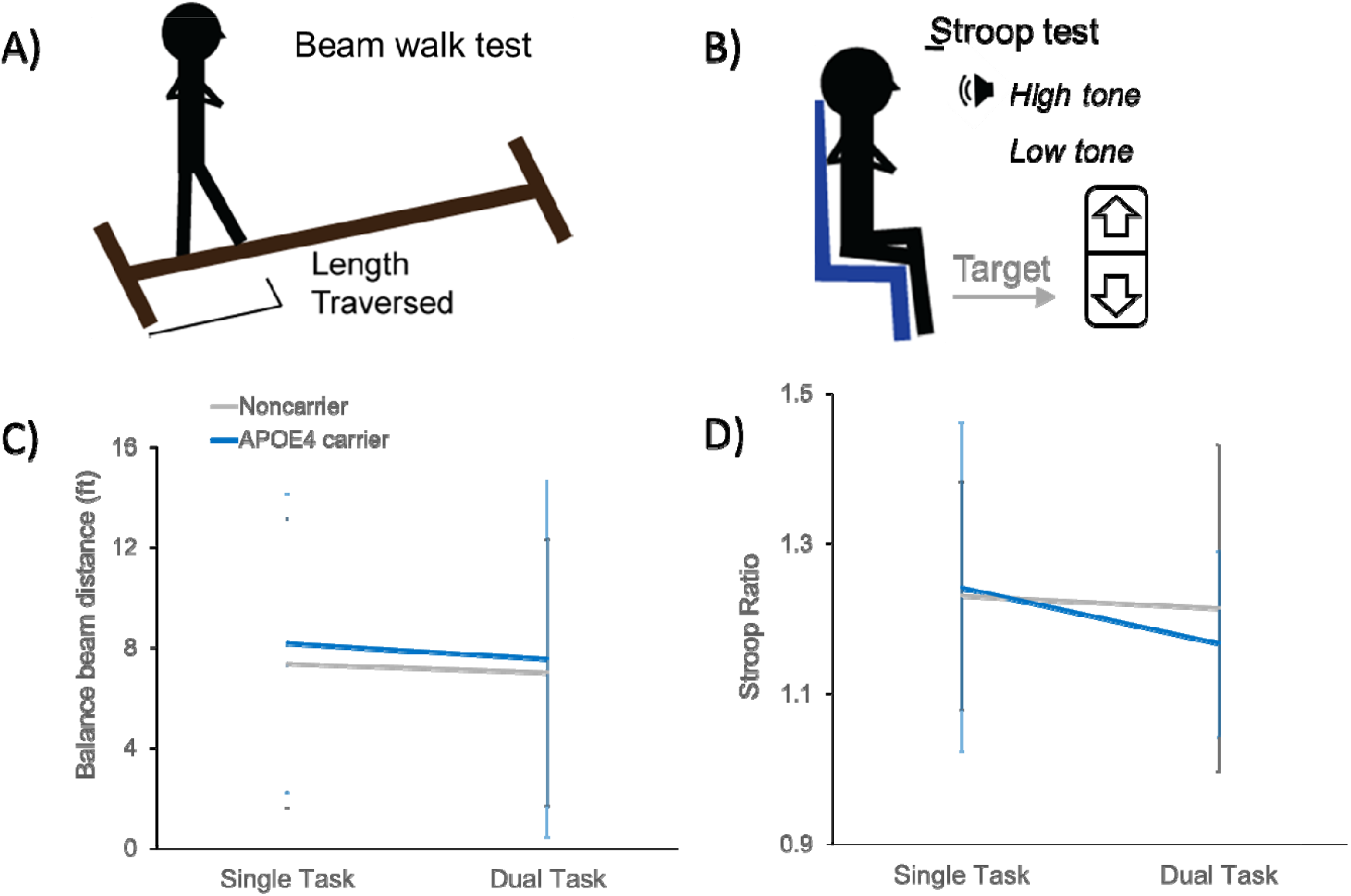
Balance and cognitive performance during single- and dual-task conditions in APOE4 carriers and noncarriers. Paradigms for assessing balance and cognitive performance across all older adults showing the beam walk task **(A)** and the Stroop response inhibition task (**B**). There were no group differences between APOE4 carriers and noncarriers on the beam walk distance traversed performance (**C**) or in cognitive Stroop task performance (**D**) during either single-or dual-task conditions. Data shown as mean ± standard deviation.

#### Dual-task (DT) balance performance

Following the single-task condition, participants completed a dual-task beam walk during simultaneous performance of a cognitive task.(53) The secondary cognitive task required the participant to verbally count backward by 3’s starting at a random integer number between 20 and 100 verbally stated by the experimenter immediately following the cue to begin the beam walk. The mean beam distance traversed and gait speed across 2 trials for each single-task and dual-task balance conditions was used to compute balance DTI (below).

#### Single-task (ST) cognitive performance

To enable more precise quantification of cognitive performance, participants completed a cognitive response inhibition task using the auditory Stroop test delivered through E-Prime Software (Pittsburgh, PA). The Stroop test is a widely used cognitive executive function test assessing the ability to inhibit the prepotent and undesired response (24,57). A 50-stimulus sequence of auditory words stating either “high” or “low” were emitted at 0.5Hz in either high or low tones, with half of the word-tone stimuli being congruent (e.g. “high” in a high tone) and half being incongruent (e.g. “high” in a low tone). Participants were asked to respond to the tone of the sound and ignore the meaning of the word in a 50-stimuli sequence delivered at 0.5Hz by clicking the up (high tone) or down (low tone) button on a hand-held clicker as quickly as possible which maintaining accuracy (**Figure 1B**). Participants completed a 10-stimuli practice trial prior to the 50-stimuli test sequence.

#### Dual-task (DT) cognitive performance

Following the single-task condition, participants completed the auditory Stroop test while simultaneously standing on an unstable balance board (Fluidstance Level Balance Board) (Santa Barbara, CA). The level of balance challenge was individually adjusted with the addition or removal of a convex board cap based on the participant’s balance ability, determined by the participant’s level of stability while standing on the board assessed by a licensed physical therapist. Participants were required to fold their arms across their chest and affix their gaze at an eye-level point on the wall while holding the remote clicker.

Stroop test accuracy was >90% for each task condition (single-task: 95% ± 6%; dual-task: 91% ± 15%), and all incorrect trials were removed from response time analyses. A Stroop ratio was computed as the mean response times of all trials with congruent stimuli over the incongruent stimuli. The Stroop ratio during each the single- and dual-task conditions was used to compute *dual-task interference (DTI)* as:

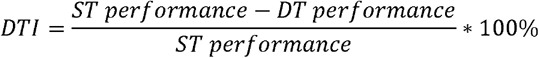

For balance DTI, the value was multiplied by (−1), such that a negative DTI value for each balance and cognition indicates worsening of performance (i.e. slower response times during incongruent relative to congruent stimuli) between the single- and dual-task conditions.(37)

### 2.4. Cerebrovascular assessment during aerobic exercise

Briefly, participants arrived to the laboratory (22 and 24 degrees Celsius) in the morning and abstained from caffeine for 12 hours, intense exercise for 24 hours, and a large meal for 2 hours. TCD was used to assess cerebral artery blood velocity (CBFv) during a single bout of moderate-intensity aerobic exercise.(49,50) A 2-MHz TCD probe (RobotoC2MD, Multigon Industries) was placed over the left temporal window. The left anterior cerebral artery (ACA) was targeted because of its vascular supply to lower extremity sensorimotor cortical regions involved in balance and walking behaviors involved in the present study. The left MCA was used if the ACA signal was absent, and the right side was used if the signal was absent on the left side. A 5-lead electrocardiogram (ECG)continuously monitored and recorded heart rhythm. Following TCD and ECG setup, we implemented a moderate-intensity aerobic exercise protocol on a recumbent stepper (NuStep T5XR) (49,50). Prior to data recording, participants familiarized themselves with the reciprocal stepping motion at a cadence of 100 steps per minute. We defined our moderate-intensity target as 45-55% of heart rate (HR) reserve using the Karvonen formula. We conducted an individualized calibration procedure to establish each participant’s optimal work resistance: beginning with the stepper set to 30 watts, we added 10 watts at regular intervals while monitoring HR until participants achieved and maintained their prescribed HR reserve zone of 45-55%. After establishing this personalized resistance level, we halted the calibration exercise and allowed participants to recover quietly for no less than 10 minutes, ensuring complete physiological stabilization before commencing the experimental trial. The experimental recording consisted of 90 seconds of seated rest on the recumbent stepper, followed by 6 minutes of continuous moderate-intensity exercise. We employed a graduated exercise initiation protocol to minimize potential Valsalva maneuvers and prevent abrupt physiological fluctuations: participants began stepping at 60% of their individualized target wattage and progressively increased resistance at 10-second intervals, achieving their full target workload 30 seconds after exercise initiation. Participants then maintained this moderate-intensity workload for the remaining duration of the 6-minute exercise period. Custom MATLAB software (The Mathworks Inc.) using an analog-to-digital data acquisition unit (NI-USB-6212, National Instruments) acquired MCAv (500 Hz), synchronized across the cardiac cycle.(49,50) Data were visually inspected and discarded when R-to-R intervals were >5 Hz or changes in peak CBFv exceeded 10 cm/s in a single cardiac cycle. Trials with <85% samples were discarded. Mean CBFv was calculated from the area under the curve for each cardiac cycle and analyses were conducted using 3-s time-binned mean values over the entire rest and exercise period, as described previously(50). We calculated cerebrovascular response (CVR) as the difference between mean CBFv during minutes 3 to 4.5 at exercise steady state and mean CBFv during the rest prior to the start of exercise.

### 2.5. *APOE* genotyping

Participants provided whole blood samples that were drawn and stored frozen at −80 degrees Celsius. Genetic analyses was performed using a Taqman single nucleotide polymorphism (SNP) allelic discrimination assay (ThermoFisher) to determine *APOE* genotype. *APOE4, APOE3*, and *APOE2* alleles were determined using Taqman probes to the two *APOE*-defining SNPs, rs429358 (C_3084793_20) and rs7412 (C_904973_10).(58,59) Individuals were classified as *APOE4* carrier in the presence of 1 or 2 *APOE4* alleles (e.g. E3/E4, E4/E4). Individuals with homozygous E3 (e.g. E3/E3) were classified as a noncarriers. We excluded any individual who carried one or two copies of *APOE2*, as *APOE2* is associated with reduced risk for Alzheimer’s disease and could affect cerebrovascular and behavioral results of the present study.(22,36)

### 2.6. Statistical analyses

We confirmed normality and heterogeneity of variance using Kolmogorov-Smirnov and Levene’s tests, respectively. We used two-way mixed analysis of variance (ANOVA) tests to assess balance beam performance and cognitive Stroop task performance between single- and dual-task conditions within each participant and between *APOE4* carriers and noncarriers. We used Pearson product-moment correlation coefficients to test relationships between CBFv response to exercise and each balance and cognitive DTI. Given the effect of *APOE4* on Alzheimer’s disease risk and the link between DTI and AD in older adults,(60–62) we then used two-way multiple linear regression (MLR) analyses (factors: APOE genotype, CBFv response to exercise, genotype-by-CBFv response to exercise) to test whether cerebrovascular-behavioral relationships differed as a function of *APOE4*. All analyses were performed using SPSS version 29 with an *a priori* level of significance set to 0.05.

## RESULTS

Two participants had unstable TCD signals with excessive noise during aerobic exercise; these participants were discarded from subsequent analysis. One participant had an ACA signal on the right side only; the right side was used for this participant. The ACA signal could be located and stabilized in 21/30participants; for the remaining 9 participants, the left MCA signal was used for analyses. Two out of these 9 participants only possessed TCD signals in the right side, and thus the right side was used. Two participants (n=1 *APOE4* carrier, n=1 noncarrier) did not follow instructions for the Stroop test during single and/or dual-task conditions; these participants were discarded from cognitive analyses. Stroop test accuracy on the single-task was 95% ± 6% and 91% ± 15% under the dual-task condition. No differences in demographics of age or sex were observed between participants in the APOE4 and noncarrier groups (*p*>0.05). There were no differences in cardiovascular or exercise metrics between groups, including blood pressure, heart rates, CBFv response to exercise and exercise wattage (see **Table 1**).

### Balance and cognitive performance under single and dual-task conditions in *APOE4* carriers compared to noncarriers

Cognitively-normal older adults with *APOE4* showed no differences in balance or cognitive performance compared to noncarriers, regardless of task condition. When comparing balance beam performance and cognitive Stroop task performance between single- and dual-task conditions as a function of *APOE4* genotype, there were no effects of condition or group. There were no group-by-condition interaction effects for metrics of balance beam distance traversed, balance beam gait speed, or cognitive Stroop ratio task performance (*F*_1,28_ ≤ 0.93, p≥0.343) and no main effects of condition (*F*_1,28_ ≤ 2.57, p≥0.120) or group (*F*_1,28_ ≤ 0.71, p≥0.406) (**Figure 1 C&D**).

### Effect of *APOE4* genotype on the relationship between cerebrovascular response to exercise and dual-task balance and cognitive performance

Older adults with *APOE4* displayed stronger cerebrovascular-behavioral relationships across balance and cognitive domains of dual-task interference compared to noncarriers. Within the balance domain, higher cerebrovascular response to exercise associated with more positive balance DTI, indicated by lesser decline in gait speed during balance beam performance under dual-task conditions in *APOE4* carriers (r=0.839, p=0.009) and showed no relationship in noncarriers (r=0.143, p=0.549) (**Figure 2A**). A trend for a similar relationship in APOE4 carriers was also observed for balance DTI on beam distance traversed in *APOE4* carriers (*r*=0.613, *p*=0.106) and showed no relationship in noncarriers (r=0.298, p=0.202), but failed to meet our adopted level of significance. Within the cognitive domain, *APOE4* carriers with a greater cerebrovascular response to exercise exhibited lesser decline in cognitive Stroop task performance from single-to dual-task conditions (r=0.832, p=0.020) while no relationship was observed in the noncarriers (r=-0.316, p=0.187) (**Figure 2B**).

**Figure 2.**
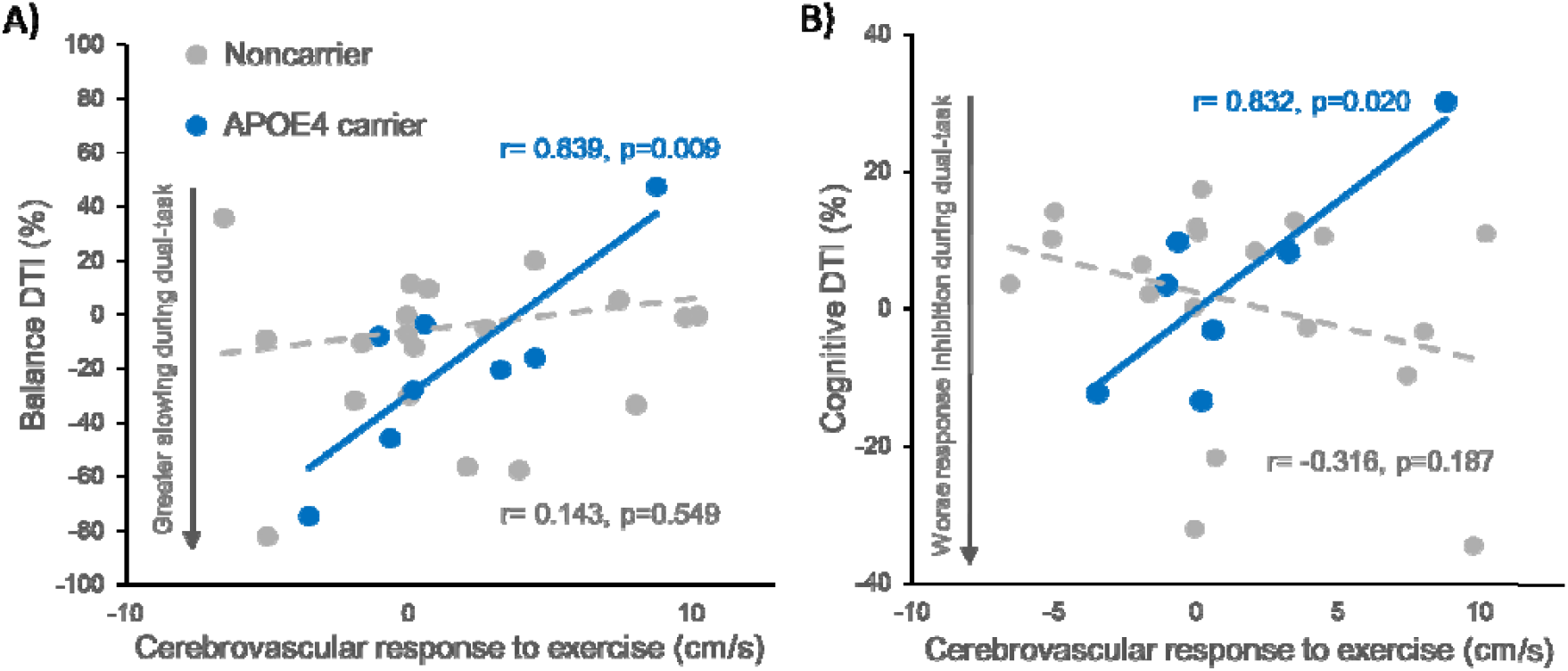
**Relationship between cerebral blood velocity (CBFv) response to aerobic exercise and dual-task interference (DTI) in APOE4 carriers and noncarriers** in each domain of balance **(A)** and cognitive performance **(B)**. There was a significant relationship in APOE4 carriers in each the balance and cognitive domains of dual-task interference, in which older adults with APOE4 who displayed higher cerebrovascular responses exhibited less balance DTI in slowing of gait speed during the beam walk task (r=0.839, *p*=0.009) (**A**) and less cognitive DTI in worsening of response inhibition performance (r=0.832, *p*=0.020) (**B**), while noncarriers showed no relationships.

We performed exploratory analyses to test whether the observed cerebrovascular-behavioral relationships significantly differed between APOE4 carriers and noncarriers. In the domain of cognitive performance, there was a significant interactive effect of *APOE4* genotype with cerebrovascular response to exercise on cognitive Stroop DTI (model: *F*_3,25_=2.42, *p*=0.093, *R*^*2*^=0.248, adjusted *R*^*2*^=0.146; interaction: t= 2.634, p=0.015), in which the relationship between higher cerebrovascular response to exercise and more positive Stroop DTI in APOE4 carriers was stronger than the relationship in noncarriers (**Figure 2B**). In the domain of balance performance, the genotype-by-cerebrovascular response to exercise interaction effect was not statistically significant (Balance beam distance traversed: model: *F*_3,27_=2.414, *p*=0.091, *R*^*2*^=0.232, adjusted *R*^*2*^=0.136; interaction: t= 1.639, p=0.114; Balance beam gait speed: model: *F*_3,27_=1.45, *p*=0.253, *R*^*2*^=0.153, adjusted *R*^*2*^=0.048; interaction: t= 1.30, p=0.208) (**Figure 2A**).

## DISCUSSION

Our study provides initial evidence for *APOE4* genotype as a biomarker for interactive effects of cerebrovascular response and clinically-relevant cognitive-balance task behavior in cognitively-normal older adults. While the role of *APOE4* in AD risk and pathology, as well as the relationships between brain vascular function and cognitive dysfunction have been well-characterized in the context of disease (e.g. MCI, AD, dementia),(20,63,64) and disease risk,(17,65) the investigation of the role of cerebrovascular function in older adults with well-preserved cognition and physical function despite advanced age (e.g. “SuperAgers”) (66,67) remains a nascent area of investigation. Here, the well-preserved cognitive-motor dual-task behavior and cerebrovascular function paired with unique cerebrovascular-behavioral relationships in this subgroup of *APOE4* carriers with an average age of 75 years provides preliminary findings for larger mechanistic studies and clinical trials to further elucidate brain resilience mechanisms in this unique aging phenotype.

In contrast to previous studies, the subgroup of *APOE4* carriers in the present study showed no difference in cerebrovascular function or cognitive-balance dual-task interference compared to noncarriers. Despite having genetically higher risk for the development of AD and advanced age, the *APOE4* carriers in the present study were cognitively-normal and showed no differences in cardio- or cerebrovascular metrics assessed here (Table 1), suggesting resilience to cognitive decline despite high genetic risk. While previous studies show that *APOE4* carriers on average display lower markers of brain health compared to noncarriers,(2,6–9) the preserved cerebrovascular response to aerobic exercise in our cohort may implicate protective, modifiable factors such as physical activity in genetic AD risk. While Whitson et al (2018) found that cognitively-normal *APOE4* carriers showed greater cognitive dual-task interference during response inhibition tasks with walking,(36) older adults in the present study were 10 years older on average (65 yo compared to 75 yo in the present cohort). Cognitive response inhibition, or the ability to suppress an undesired default or automatic response in the presence of interfering stimuli, has been identified as an early marker of impaired cognition and can distinguish between older adults who are cognitively normal compared to those diagnosed with mild cognitive impairment (24). Existing evidence shows that brain resilience, the ability to cope with challenge, declines with age and influences the susceptibility to dementia.^1^ Simultaneous cognitive loading during balance control may probe neurocognitive and neuromotor resilience to physiologic challenge.(35) While the precise neural mechanisms underpinning the interplay between cognition and motor control is unclear, one theory poses that dual-task performance is limited by an individuals’ neural capacity that ultimately limits behavioral task complexity.(31–33) Because preserved function becomes less prevalent with advanced age (47), our preliminary results here identify cognitively-normal *APOE4* carriers in advanced age and who maintain normal cognition and measures of brain vascular function as a potential model of aging resilience. The absence of group differences in cerebrovascular function may implicate protective, modifiable factors such as physical activity, whose multifaceted effects (68) could support both the cerebrovascular response to exercise and simultaneous cognitive-motor task demands during dual-tasking in this subgroup with high genetic AD risk.

Findings of the present study provide initial evidence for a neuroprotective role of cerebrovascular function in this vulnerable subgroup of APOE4 carriers that extends beyond single-task cognitive performance. Cerebrovascular-behavioral relationships in APOE4 carriers in the present study may be consistent with other recent findings from our group in a larger cohort (n=47 *APOE4* carriers), showing tha*t APOE4* carriers appear to display greater cortically-mediated anticipatory increases in cerebral blood velocity immediately prior to initiating whole-body movement, implicating these differences in cerebrovascular function as a potential compensation for other impaired cerebrovascular metrics identified in these individual older adults.(18) Notably, *APOE4* carriers diagnosed with MCI or AD did not display anticipatory increases in cerebral blood velocity with movement, implicating a potential neuroprotective role of compensatory cerebrovascular function for disease.(18) These preliminary findings may also generally support the vascular hypothesis of AD, which points to cardiovascular disease and cerebral hypoperfusion as origins in the inflammatory cascade precipitating the development of AD.(10,69)

In the face of competing attentional demand, the present findings provide preliminary evidence that maintaining higher cerebrovascular function may support older adults’ neural capacity for dual cognitive-balance task performance. Unique relationships in *APOE4* carriers between cerebrovascular response and cognitive-balance performance under dual-task conditions could imply an increased role of cerebrovascular health for behaviors that necessitate greater cortical resources in this older adult subgroup. While the precise neural mechanisms underpinning the interplay between cognition and motor control is unclear, one theory poses that dual-task performance is limited by an individuals’ neural capacity that ultimately limits behavioral task complexity.(31–33) Building upon our own work (23) and other prior studies involving single-task conditions,(70–72) our results support the notion that brain resilience and functional capacity are probed under challenging behavioral conditions(73) such as cognitive-balance dual-tsking, and that higher cerebrovascular health may promote greater neurocognitive and neuromotor resilience to this physiologic challenge in older adults with high genetic vulnerability to AD. These preliminary results identify cerebrovascular health as a promising target for interventions aimed at maximizing neurocognitive and neuromotor resilience linked to age-related disease, supporting a precision-based approach with individual genetic risk.

### Strengths and limitations

Subjecting participants to a comprehensive cognitive battery enabled us to confidently capture cognitively-normal older adults in advanced aging, who may be underrepresented in cognitively-normal cohorts given the pivotal role age plays in cognitive decline.(47) Nevertheless, the modest sample size and cross-sectional nature identifies areas for further investigation by larger investigations with longitudinal measures. While there were no differences between age and sex between *APOE4* carriers and noncarriers, age and sex affect cerebrovascular-behavioral relationships(46,51) and warrants further investigation in larger studies. While transcranial Doppler ultrasound offers clinical applicability, it offers limited regional specificity compared to MR-based techniques.(74) The focus on a single cognitive domain across balance dual-task conditions restricts interpretation to cognitive response inhibition contexts. Variations may arise when applying this paradigm to other cognitive domains (e.g., working memory).(36) The generalizability and reproducibility of our findings are constrained by the underrepresentation of non-white races as a result of recruitment in these segments of community-dwelling older adults, emphasizing the necessity for increased outreach efforts.

## CONCLUSION

The present findings provide preliminary evidence for cerebrovascular function as a mechanism supporting neuromotor resilience to cognitive loading under challenging balance conditions in older adult APOE4 carriers who are resistant to cognitive decline. These findings extend the clinical relevance of *APOE4* as a genetic biomarker beyond the context of cognitive function and implicate its relevance in cortically-mediated balance control in motor behavioral contexts with aging. Our findings from a model of healthy aging offer the potential to create a paradigm shift in the clinical and scientific framework of balance control with aging, suggesting that physical activity interventions targeting increased cerebrovascular function may offer an effective method to delay or prevent early clinical manifestations of cognitive interference in balance control in older adult subgroups with high genetic vulnerability to AD.

## Author contributions

JP conceived of the present research question, designed this study, collected and analyzed the data. JP and SB obtained funding to support this study. SB and the KU Alzheimer’s Disease Research Center contributed to recruitment. JP constructed the figures and the first draft of this manuscript. SB contributed the analysis tools and expertise for interpreting cerebrovascular data. All authors discussed the results and contributed to the final manuscript.

We would like to acknowledge Emily Hazen for her contribution to data collection and Dr. Jill Morris for her contribution to APOE genotyping.

